# A mycovirus identified from an outbreak fungal species complex reveals a parallel antiviral system

**DOI:** 10.64898/2026.09.15.751652

**Authors:** Zhenghao Shen, Zeyu Duan, Xiaoyu Ma, Linhao Song, Xin Luo, Jialong Xu, Ruiqiao Li, Yinglu Cui, Xiao Liu, Linqi Wang

## Abstract

Mycoviruses are prevalent across major fungal lineages, yet none have been reported in the *Cryptococcus gattii* species complex (CGSC), the causative agent of the largest known fungal outbreak in immunocompetent hosts. Consequently, the occurrence of mycoviruses and the host determinants governing their infection in this important pathogenic clade remain unexplored. Through screening of 386 *Cryptococcus* isolates, we identified a mycovirus, designated CgTV1, in an environmental isolate of the CGSC. CgTV1 is a double-stranded RNA (dsRNA) mycovirus of the genus *Totivirus*, with a genome encoding a capsid protein and an RNA-dependent RNA polymerase. We showed that CgTV1 establishes a cryptic infection, because neither viral presence nor its clearance significantly alters host transcriptomic profiles, suggesting the existence of mechanisms that maintain virus–host homeostasis. Population genomic analyses reveal that an intact RNAi pathway constitutes a critical genetic barrier, and CgTV1 persistence strictly requires loss-of-function mutations in both *AGO1* and *AGO2*. By pan-transcriptomics across 43 CGSC strains, we further uncovered an RNase III gene, *CAI1*, which exhibits high expression in the mycovirus-positive isolate. Biochemical analysis indicates that Cai1 directly recognizes and degrades CgTV1 dsRNA, thereby constraining excessive mycoviral proliferation. We further demonstrate that Dicer and Cai1 act in parallel to maintain virus–host homeostasis: their combined ablation results in higher viral loads and more pronounced changes in host gene expression than ablation of either factor alone. Collectively, our findings provide the evidence of the occurrence of a mycovirus in the CGSC and reveal parallel mechanisms that enable cryptic mycoviral infections.

## Introduction

Mycoviruses are viruses that infect fungi and replicate stably within fungal cells (1, 2). Owing to the lack of an extracellular transmission phase, their life cycle is entirely dependent on the host cell (2, 3). Based on genome type, mycoviruses are classified into double-stranded RNA (dsRNA), positive- and negative-sense single-stranded RNA (ssRNA), and single-stranded DNA (ssDNA) viruses, with dsRNA and ssRNA viruses being the most prevalent (4). The majority of mycoviruses establish cryptic or asymptomatic infections (3, 5, 6), whereas only a minority have been reported to confer detectable phenotypic alterations, such as changes in host growth, secondary metabolism, or virulence (7–10). Extensive research has elucidated the molecular mechanisms underlying virus–host interactions in plant-pathogenic fungi (9, 11, 12). In contrast, both the prevalence of mycoviruses and the host–virus dynamics in human-pathogenic fungi remain largely uncharacterized, with such interactions documented to date in only a few lineages (13–16).

Given that antiviral systems in microbes share evolutionary homology with key components of human antiviral immunity, fungi serve as valuable model organisms for studying viral defense mechanisms in eukaryotes. As a classical antiviral pathway in eukaryotes, the RNA interference (RNAi) pathway has been shown to suppress viral replication in various fungi (17, 18). For example, the key components of RNAi, including Dcr1, Dcr2, and Ago2, have been reported to be involved in suppressing mycovirus replication across fungal species (17, 19, 20). More recently, the mitochondrial DNA/RNA endonuclease Nuc1 was identified in *Saccharomyces cerevisiae* as a novel anti-mycoviral factor, and its mammalian homolog promotes genome fragmentation during virus-induced programmed cell death (21, 22). Despite these important advances, the understanding of the diversity and synergistic interplay of fungal antiviral strategies remains very limited.

The genus *Cryptococcus* includes two human pathogen clades: *Cryptococcus neoformans* species complex (CNSC) and the *Cryptococcus gattii* species complex (CGSC) (23, 24). These fungi are widely distributed in the environment and are designated by the WHO as major human fungal pathogens (25, 26). The CNSC accounts for the majority of cryptococcosis cases worldwide, whereas the CGSC can infect immunocompetent individuals and has caused the largest known outbreak in this population among fungal pathogens (27, 28). Despite the high clinical significance of the CNSC and CGSC, no mycoviruses have yet been reported in these two pathogenic clades.

To determine whether mycoviruses exist in the CNSC and CGSC, we screened 386 *Cryptococcus* isolates and identified a dsRNA virus in the CGSC, which we designated *Cryptococcus gattii* totivirus 1 (CgTV1). By integrating population comparative genomics, transcriptomics, and molecular genetics, we revealed that the mycovirus-positive strain harbors loss-of-function mutations in both Argonaute genes, *AGO1* and *AGO2*, and that reintroduction of a functional copy of either gene effectively suppresses viral persistence *in vivo*. In parallel, we identified an endoribonuclease, Cai1 (<u>C</u>ryptococcal <u>A</u>ntiviral Immunity Protein 1), belonging to the RNase III family, which functions as an important antiviral factor and acts synergistically with Dicer to restrict mycoviral loads and maintain host transcriptomic homeostasis independently of Argonaute proteins. In conclusion, this study fills the phylogenetic gap for mycoviruses in *Cryptococcus* and elucidates the synergistic interplay among different antiviral systems that enables cryptic infection of CgTV1 in its host.

## Results

### Identification of a dsRNA virus in the CGSC

We investigated mycovirus infection across the CNSC and CGSC by screening 386 clinical and environmental isolates, encompassing *C. neoformans, C. deneoformans, C. gattii, C. deuterogattii, C. bacillisporus*, and *C. tetragattii* (Dataset S1) via metavirome sequencing. Following quality filtering, *de novo* assembly, and BLAST analysis, a single 4,558-bp contig was identified, which is similar to *Scheffersomyces segobiensis* virus L (SsVL), a dsRNA virus belonging to the genus *Totivirus* (Table S1). These *in silico* predictions were validated using RT-PCR with specific primers targeting the viral contig sequence. Among all screened *Cryptococcus* isolates, only the *C. gattii* CG1 yielded a specific amplicon band (Fig. S1A). Total nucleic acids were extracted from *C. gattii* CG1 according to an established protocol (29). A distinct nucleic acid band of approximately 4.5 kb was observed in the *C. gattii* CG1 strain and subsequently subjected to digestion treatments with DNase I, S1 nuclease, and RNase A (Fig. 1A). The results showed that this band was a dsRNA molecule. By employing rapid amplification of cDNA ends (RACE) coupled with Sanger sequencing, the initial contig was further extended to yield a complete viral genome of 4,569 bp (Fig. 1B). The genome comprises two open reading frames (ORFs), encoding a 78.28 kDa capsid protein (CP) and an 84.90 kDa RNA-dependent RNA polymerase (RdRp), respectively (Fig. 1B). Maximum likelihood phylogenetic analysis based on the CP and RdRp amino acid sequences placed this mycovirus within the genus *Totivirus* of the family *Orthototiviridae* (Fig. 1C). Pairwise alignment revealed that the CP and RdRp proteins shared only 13.84%–39.12% and 27.20%–52.94% amino acid identity, respectively, with their closest homologs in the genus (Fig. 1D). This low sequence similarity indicates that the virus represents a new species within the genus *Totivirus* (30).

**Figure 1.**
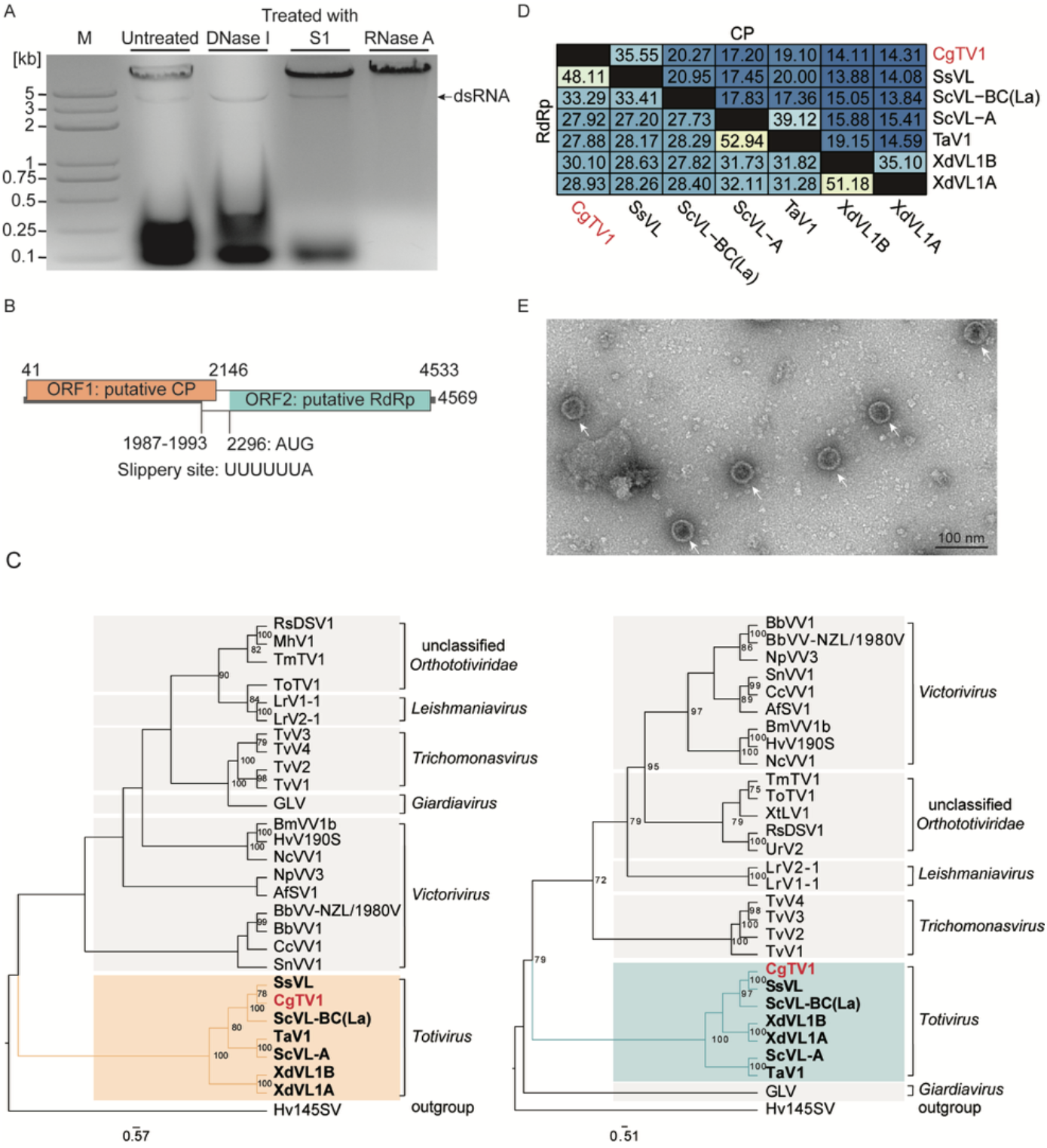
Identification and characterization of the mycovirus CgTV1 in *C. gattii*. (A) Agarose gel electrophoresis profile of total nucleic acids extracted from *C. gattii* CG1. Samples were either left untreated or digested with DNase I, S1 nuclease, or RNase A. The black arrow indicates the viral dsRNA band. M, DNA molecular weight marker. (B) Genomic organization of CgTV1. The full-length genome is 4,569 bp. ORF1 (nt 41-2,146) encodes a putative 78.28 kDa capsid protein. AUG (nt 2,296-2,298), the start codon in ORF2, is downstream of the ORF1 stop codon. ORF2 (nt 1,986-4,533) encodes a putative 84.90 kDa RNA-dependent RNA polymerase. ORF2 should be translated as a fusion protein with ORF1 through a −1 ribosomal frameshift, the slippery site UUUUUUA (nt 1,987–1,993) was found in the overlapping regions. (C) Maximum likelihood phylogenetic analysis of CP (left) and RdRp (right) sequences using IQ-TREE. Numbers at the nodes represent bootstrap percentages. The position of CgTV1 is indicated in red. Scale bars represent the number of amino acid substitutions per site. Virus abbreviations are detailed in Table S4 and Table S5. (D) Amino acid sequence identity matrix for CgTV1 and related species within the genus *Totivirus*. The upper right triangle displays CP identities, while the lower left triangle displays RdRp identities (values are in percentages). (E) Representative TEM image of VLPs purified from *C. gattii* CG1. White arrows indicate VLPs. Scale bar, 100 nm. An additional representative TEM image is provided in Figure S2.

To further confirm the presence of the mycovirus, we isolated and purified virus-like particles (VLPs) using sucrose density gradient centrifugation (31). Nucleic acid detection showed that the viral genome was predominantly detected in the 50%–60% sucrose fractions (Fig. S2A). Transmission electron microscopy (TEM) confirmed the presence of abundant VLPs approximately 40 nm in diameter in these fractions (Fig. 1E and Fig. S2B, C). These morphological characteristics are consistent with those of previously reported members of the genus *Totivirus* (32). Collectively, we identified a dsRNA mycovirus in *C. gattii* and designated it as *Cryptococcus gattii* totivirus 1 (CgTV1), whose complete genome sequence has been deposited in GenBank under accession number PZ721968.

### CgTV1 persistence strictly requires loss-of-function mutations in both *AGO1* and *AGO2*

Given the extremely low prevalence of mycoviruses among *Cryptococcus* isolates (< 0.3%), we hypothesized that the critical antiviral components in this virus-positive strain might be compromised by loss-of-function mutations. We tested this hypothesis using our previously published genomic data from 43 *C. gattii* strains (33), a subset of the above-screened 386 isolates, which included the CgTV1-positive CG1 strain and 42 negative strains. To identify the genetic determinants underlying persistent mycovirus infection, we screened for genetic mutations harbored by the *C. gattii* CG1 strain that occurred at low frequencies among mycovirus-negative strains. Compared with the reference strain *C. gattii* WM276 (34), *C. gattii* CG1 harbored 693 high-impact mutations, collectively affecting 369 genes (Fig. 2A and Dataset S2). Among these genes, we identified concurrent loss-of-function mutations in the key RNAi components *AGO1* and *AGO2* in *C. gattii* CG1. These mutations occured at low population frequencies of 4.7% and 20.9%, respectively (Fig. 2B). Specifically, in *C. gattii* CG1, *AGO1* harbored a splice-donor mutation (c.789+2T>C) and a nonsense mutation that truncates the protein within the PAZ domain (c.991C>T, p.Arg331*). Meanwhile, *AGO2* contained a frameshift mutation in the PAZ domain (c.996dupT, p.Ala333fs), a splice donor mutation (c.1198+2C>T), and a 3’ terminal deletion encompassing the Piwi domain (c.2668-9_*12del). Sanger sequencing confirmed the presence of these mutations, consistent with the genomic analysis (Fig. 2C, D).

**Figure 2.**
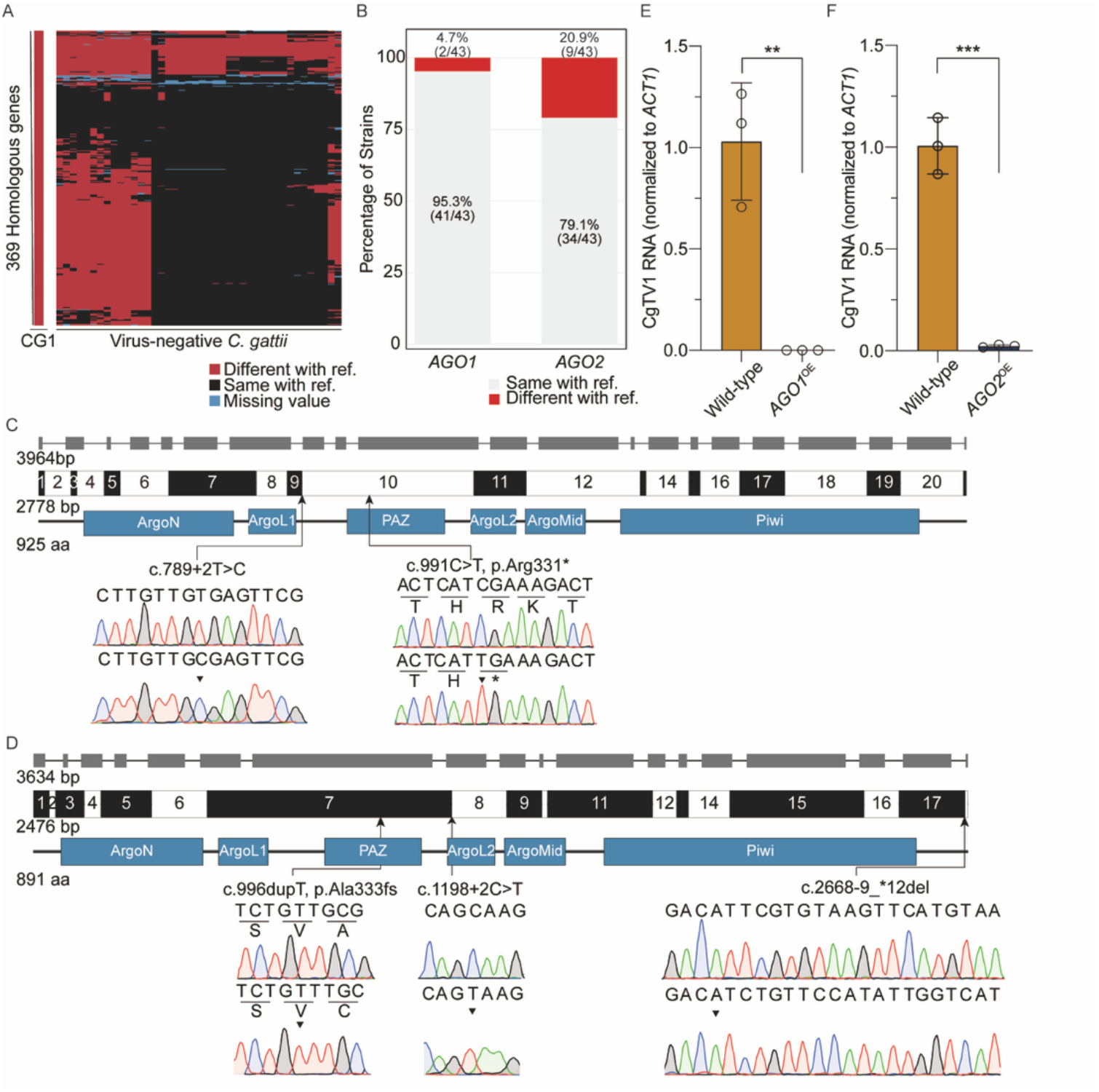
Population comparative genomic analysis and functional validation of *AGO1* and *AGO2* in *C. gattii*. (A) Heatmap of genes harboring high-impact variations across 43 *C. gattii* strains. Red, sites differing from the reference genome *C. gattii* WM276; black indicates identical sites; blue indicates missing sequencing data. (B) Population distribution of *AGO1* and *AGO2* genotypes across 43 *C. gattii* isolates. Red represents putative loss-of-function variants (including the alleles identified in *C. gattii* CG1); gray represents functional alleles matching the reference strain WM276. (C, D) Gene structures, cDNA exon arrangements, protein domain compositions, and mutation profiling of *AGO1* (C) and *AGO2* (D). For each panel, schematic diagrams (from top to bottom) depict the genomic gene structures (grey boxes represent exons), cDNA exon arrangements (alternating black and white boxes with exon numbers), and protein domains (blue boxes). Representative Sanger sequencing chromatograms display the identified mutations in *C. gattii* CG1 (bottom traces) aligned against *C. gattii* WM276 controls (top traces). Mutated nucleotides are indicated by arrow heads, with arrows marking their locations relative to exons and protein domains. Domain abbreviations: ArgoN, Argonaute N-terminal domain; ArgoL1, Argonaute linker 1 domain; PAZ, PAZ domain; ArgoL2, Argonaute linker 2 domain; ArgoMid, Argonaute Mid domain; Piwi, Piwi domain. (E, F) Relative viral RNA level in *AGO1* (E) and *AGO2* (F) overexpression strains was measured by RT-qPCR. Mycoviral RNA levels were normalized to the host *ACT1* gene and calculated relative to the wild-type strain. Open circles represent individual biological replicates (n = 3), and error bars indicate mean ± SD. \*\**P* < 0.01, \*\*\**P* < 0.001 (two-tailed, unpaired Student’s *t*-test).

Assessing whether functionally intact RNAi machinery can suppress viral accumulation involved overexpressing the full-length, functional *AGO1* or *AGO2* genes from *C. gattii* WM276 (35) in the *C. gattii* CG1 strain. RT-qPCR revealed that overexpression of either *AGO1* or *AGO2* significantly reduced viral RNA levels (Fig. 2E and F). Collectively, these findings indicate that the loss-of-function mutations in both *AGO1* and *AGO2* are indispensable for CgTV1 persistence in *C. gattii*.

### Population transcriptomics identifies an endoribonuclease as an antiviral factor

To determine whether CgTV1 presence affects the host transcriptional response, we treated the *C. gattii* CG1 strain with 2′-C-methylcytidine (2CMC) and confirmed successful viral clearance by RT-PCR (Fig. S1B). Comparative transcriptomics (RNA-seq) between the wild-type and its virus-cured derivative revealed no differentially expressed genes (DEGs), indicating that the host maintains transcriptional homeostasis in the presence of CgTV1(Fig. S3). To further identify host genes responsible for this homeostasis, we analyzed the previously published transcriptomic data from the 43 *C. gattii* strains and performed a comparative transcriptomic analysis (33). We hypothesized that host antiviral factors would be highly expressed in the CgTV1-positive strain compared with the virus-negative strains, and therefore applied a two-step filtering strategy to the population transcriptomic data. First, we identified genes in the virus-positive strain *C. gattii* CG1 that ranked in the top 10% of population expression levels and exhibited statistically significant divergence (*P* < 0.05; Fig. 3A and Dataset S3). Second, to ensure candidate genes exhibit robust transcriptional activity, we calculated Robust Z-score for all genes in *C. gattii* CG1, retaining only those exceeding the internal median expression level of the *C. gattii* CG1 transcriptome (Z > 0; Fig. 3A and Dataset S3). This two-step screening yielded a final set of 532 candidate genes for downstream analysis (Dataset S3). Among these genes, we focused on those with predicted dsRNA nuclease activity, as such enzymes have been reported in diverse eukaryotes to directly restrict dsRNA virus infections by degrading viral dsRNA. Among all candidates harboring dsRNA nuclease domains, *FUN_002417* ranked the highest. This gene encodes a predicted RNase III family endoribonuclease, which we designate *CAI1* (<u>C</u>ryptococcal <u>A</u>ntiviral <u>I</u>mmunity Protein 1). Pfam-based prediction and complex structure prediction collectively confirmed that Cai1 possesses a canonical RNase III domain architecture (RIIID and dsRBD) and may bind dsRNA (Fig. 3B), supporting its potential to directly recognize and cleave mycoviral dsRNA.

**Figure 3.**
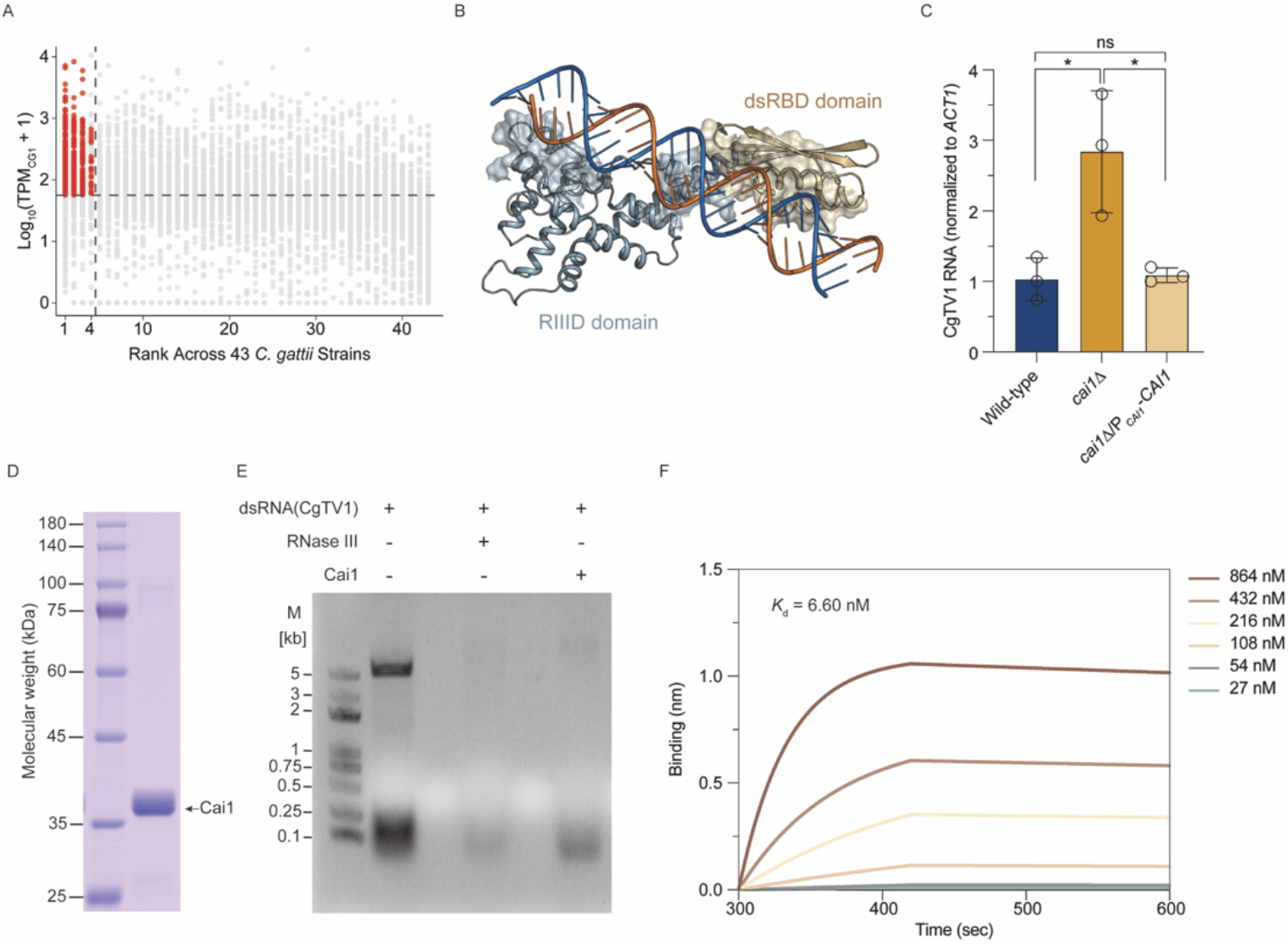
Discovery and biochemical characterization of the antiviral endoribonuclease Cai1. (A) Candidate gene screening based on transcript abundance and population expression rank. The horizontal axis represents the expression rank across 43 *C. gattii* strains, and the vertical axis represents the transcript abundance log_10_(TPM_CG1_+1). Red dots highlight prioritized candidate genes meeting the screening criteria: population-level specificity (Rank ≤ 4, top 10% across the population, vertical dashed line), significant expression divergence (Z-score > 1.64) and substantial expression in *C. gattii* CG1 (Robust Z > 0, above the transcriptome median; horizontal dashed line). Gray dots denote background genes. (B) Complex structure prediction illustrates the binding interaction between Cai1 and dsRNA. Cai1 is shown in ribbon and surface representations, highlighting the RNase III catalytic domain (RIIID, gray-blue) and the double-stranded RNA-binding domain (dsRBD, tan) engaging a 33-bp dsRNA duplex (blue and orange strands). (C) Relative viral RNA level in wild-type, *cai1*Δ, and *cai1*Δ/P_*CAI1*_-*CAI1* strains were measured by RT-qPCR. Mycoviral RNA levels were normalized to the host *ACT1* gene. Open circles represent individual biological replicates (n = 3), and error bars indicate mean ± SD. \**P* < 0.05, \*\**P* < 0.01; ns, not significant (one-way ANOVA). (D) SDS-PAGE gel of recombinant Cai1 protein used for downstream assays. The left lane contains the molecular weight marker (kDa). (E) *In vitro* enzymatic cleavage assay of native CgTV1 dsRNA. Recombinant Cai1 or commercial RNase III (positive control) was incubated with CgTV1 dsRNA as indicated (+/-). M, DNA molecular weight marker. (F) Bio-layer interferometry (BLI) sensorgrams detailing the binding kinetics between Cai1 protein and native CgTV1 dsRNA. Curves representing variable Cai1 protein concentrations (27 nM to 864 nM) are shown, with the calculated equilibrium dissociation constant (*K*_d_) indicated.

We evaluated the role of *CAI1* in host antiviral defense by generating a *CAI1* deletion strain (*cai1*Δ) using the CRISPR-Cas9 system (35), as well as a genetically complemented strain (*cai1*Δ/P_*CAI1*_-*CAI1*). RT-qPCR analysis revealed that the CgTV1 viral loads were significantly elevated in the *cai1*Δ mutant compared to the parental strain (Fig. 3C). Genetic complementation restored viral accumulation to parental levels (Fig. 3C), demonstrating that *CAI1* plays a crucial role in restricting mycovirus replication *in vivo*. We next assessed whether Cai1 possesses dsRNA-specific cleavage activity. Accordingly, we heterologously expressed and purified the recombinant protein in *Escherichia coli* (Fig. 3D). *In vitro* enzymatic assays were conducted by incubating purified Cai1 with synthetic 33-bp ssRNA, ssDNA, dsDNA, and dsRNA (36), as well as native CgTV1 dsRNA. Cai1 displayed robust cleavage activity toward both synthetic and native CgTV1 dsRNA, while exhibiting no detectable activity toward ssRNA, ssDNA, or dsDNA (Fig. 3E and Fig. S4A). To further characterize the binding affinity of Cai1 for native CgTV1 dsRNA, we quantitatively measured their interaction using Bio-Layer Interferometry (BLI). Cai1 exhibited high binding affinity for native CgTV1 dsRNA, with an equilibrium dissociation constant (*K*_d_) of 6.60 nM (Fig. 3F). Consistently, biotinylated dsRNA pull-down assays demonstrated that CgTV1 dsRNA successfully captured and enriched Cai1 (Fig. S4B). Collectively, these findings establish Cai1 as an important antiviral factor required for constraining mycoviral loads in *C. gattii*.

### Dicer and *CAI1* act in parallel to maintain transcriptomic homeostasis in *C. gattii* CG1

Given that Dicer and Cai1 both belong to the RNase III family, we further investigated whether they act complementarily or redundantly. In *C. gattii* CG1, the two Dicer genes, *DCR1* and *DCR2*, are arranged in tandem. We therefore generated the *dcr1*Δ*dcr2*Δ double mutant and the *cai1*Δ*dcr1*Δ*dcr2*Δ triple-knockout strains. RT-qPCR analysis revealed that viral accumulation was significantly elevated in *dcr1*Δ*dcr2*Δ double mutant compared to the parental strain, which is deficient in both *AGO1* and *AGO2* (Fig. 4A). This result suggests that Dicer functions independently of Argonaute to suppress mycovirus replication in *C. gattii* CG1.

**Figure 4.**
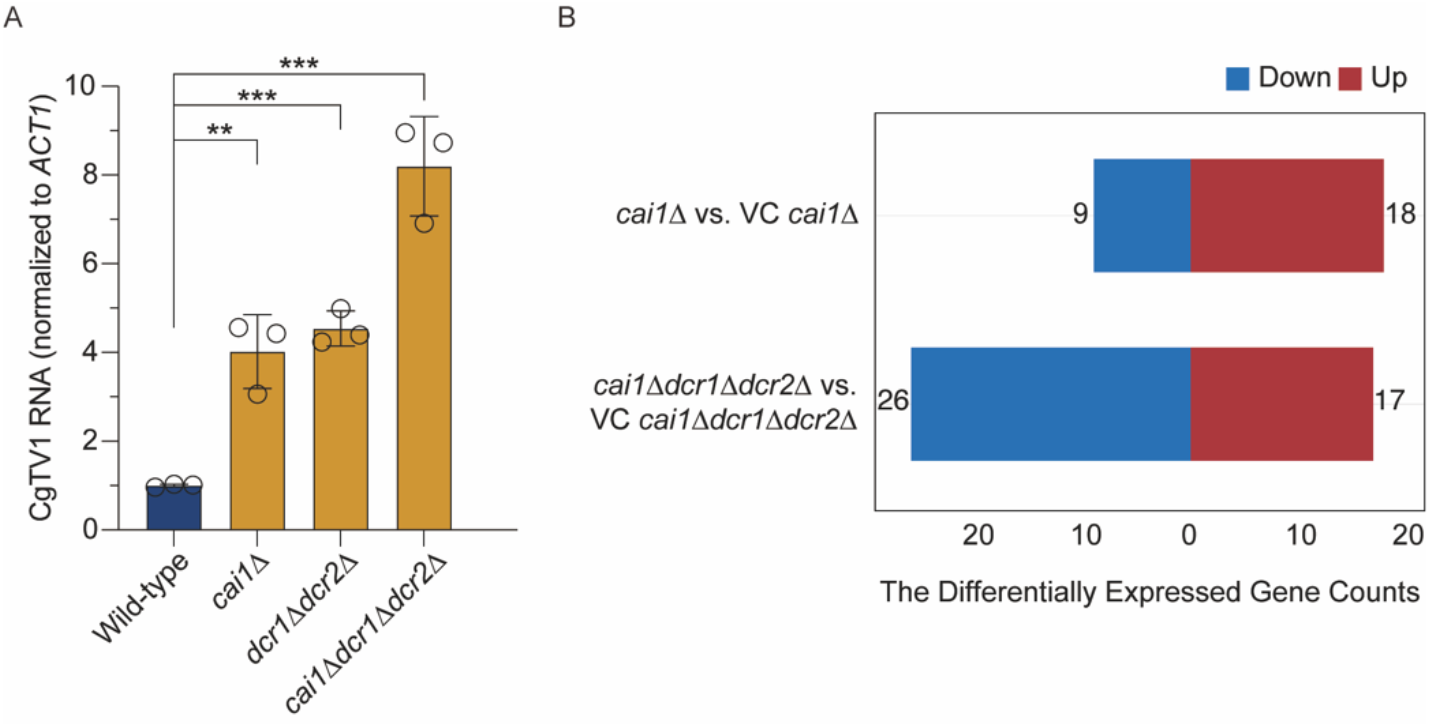
The interplay between CgTV1 and *C. gattii* endoribonucleases and RNAi system. (A) Relative viral RNA level in wild-type, *cai1*Δ, double-knockout (*dcr1*Δ*dcr2*Δ) and triple-knockout (*cai1*Δ*dcr1*Δ*dcr2*Δ) strains measured by RT-qPCR. Mycoviral RNA levels were normalized to the host *ACT1* gene and calculated relative to the wild-type strain. Open circles represent individual biological replicates (n = 3), and error bars indicate mean ± SD. \*\**P* < 0.01, \*\*\**P* < 0.001, \*\*\*\**P* < 0.0001 (one-way ANOVA). (B) Differentially expressed gene counts across mutant strains relative to their corresponding virus-cured controls. Red and blue bars indicate significantly up-regulated (*P*_adj_ < 0.05, log_2_(fold change) > 0.6) and down-regulated (*P*_adj_ < 0.05, log_2_(fold change) < −0.6) genes.

We further showed that the triple-knockout strain (*cai1*Δ*dcr1*Δ*dcr2*Δ) exhibited even higher viral loads than either the *cai1*Δ single mutant or the *dcr1*Δ*dcr2*Δ double mutant (Fig. 4A). Together, these data demonstrate that the RNase III endoribonucleases Cai1 and Dicer function independently of Argonaute, acting as a parallel antiviral mechanism that complements canonical RNAi to form a comprehensive antiviral defense network in *C. gattii* CG1 (Fig. 5). Consistent with this, comparative transcriptomics identified 27 and 43 DEGs in the *cai1*Δ and *cai1*Δ*dcr1*Δ*dcr2*Δ mutants, respectively, relative to their respective virus-cured counterparts (Fig. 4B, Dataset S4 and Dataset S5). These results indicate that, in the presence of the mycovirus, Cai1 and Dicer jointly contribute to the maintenance of host transcriptomic homeostasis.

**Figure 5.**
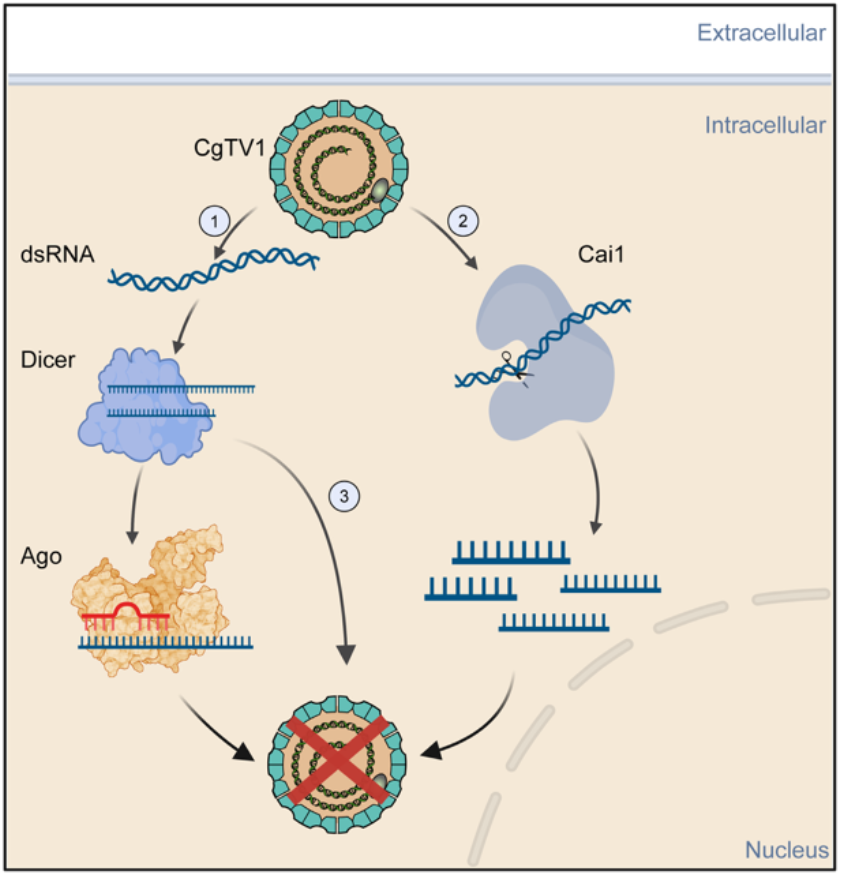
Proposed model of the parallel RNAi system and endoribonucleases in suppressing CgTV1 accumulation. Within *C. gattii* cells, CgTV1 infection exposes viral double-stranded RNA (dsRNA), triggering a multilevel antiviral response in the host. Mechanism ① represents the classical RNAi pathway, wherein viral dsRNA is recognized by Dicer and cleaved into small interfering RNAs (siRNAs). These siRNAs are subsequently loaded into the Argonaute (Ago) protein complex to mediate sequence-specific RNA silencing, thereby inhibiting viral replication. Mechanism ② outlines the Cai1-mediated degradation mechanism, in which the host restriction factor Cai1 directly recognizes and specifically cleaves viral dsRNA. Mechanism ③ illustrates an Ago-independent, Dicer-mediated direct restriction. Upon recognition of viral dsRNA, Dicer directly restricts viral assembly and replication without relying on the downstream Ago-mediated RISC. The virion illustration was adapted from ViralZone. This figure was created with Biorender.com.

## Discussion

We systematically screened 386 environmental and clinical *Cryptococcus* isolates and identified a dsRNA mycovirus in *C. gattii*, designated CgTV1 (Fig. 1A and Fig. S1A). Genomic characterization places CgTV1 within the genus *Totivirus* (Fig. 1C, Table S4 and Table S5), with its dsRNA genome encoding a CP and an RdRp, the latter translated via a −1 ribosomal frameshift (Fig. 1B). TEM analysis further confirmed the presence of virus-like particles with dimensions consistent with those of previously reported totiviruses (Fig. 1E and Fig. S2B, C) (32). Our findings thus expand the recognized host range of mycoviruses to include the *Cryptococcus* pathogenic species complex, a major group of human fungal pathogens.

Compared with the relatively high prevalence of RNA mycoviruses in some species, such as *S. cerevisiae* (~22–51%)(37, 38), the natural prevalence in *Cryptococcus* isolates is exceedingly low (1/386, ~0.26%). This observation is consistent with a recent preprint that, through transcriptome-based mining of 2,940 *C. neoformans* RNA-seq datasets, identified three isolates carrying a totivirus (39). These results suggest that the *Cryptococcus* pathogenic species complex may possess multilayered antiviral systems that impose a high barrier to mycovirus prevalence. In *Cryptococcus*, the RNAi pathway may serve as an important antiviral defense mechanism. Unlike in *S. cerevisiae*, the RNAi machinery is universally conserved across *Cryptococcus* species, with *C. deuterogattii* being the sole naturally deficient exception (40). Our study found that loss-of-function mutations in both Argonaute genes, *AGO1* and *AGO2*, were critical for CgTV1 persistence, and reintroduction of functional copies of either gene significantly restricted viral abundance (Fig. 2).

Transcriptomic analysis revealed that CgTV1 carriage did not significantly alter host transcription (Fig. S3), suggesting the existence of additional antiviral strategies that maintain virus–host immune homeostasis. We found that Dicer, another key component of the RNAi pathway, plays an independent antiviral role. In the *AGO*-defective *C. gattii* CG1 strain, simultaneous deletion of *DCR1* and *DCR2* significantly elevated viral loads (Fig. 4A), indicating that Dicer functions independently of Argonaute. These results imply that *DCR1* and *DCR2* may be under selective pressure to maintain CgTV1 homeostasis, as they remained unmutated in the virus-positive strain. Using population transcriptomics, we further identified another RNase III family member, Cai1. Through biochemical, genetic, and transcriptomic analyses, we demonstrated that Cai1 directly binds and efficiently cleaves CgTV1 dsRNA (Fig. 3F, G). Deletion of *CAI1* in the virus-positive background increased viral loads (Fig. 3D) and led to more pronounced transcriptomic changes compared with the virus-cured counterpart (Fig. 4B). These results establish an antiviral role for Cai1 in *C. gattii* CG1. Moreover, Cai1 and Dicer act in parallel as the combined deletion of both resulted in even higher viral loads than either single deletion (Fig. 4A). The discovery of both Cai1 and Dicer in antiviral defense suggests that the RNase III family may serve as an important antiviral protein family in fungi (Fig. 5).

Phenotypic analyses showed that, despite significantly elevated CgTV1 loads, the mutant strains did not exhibit obvious growth changes under 40 tested growth conditions (Fig. S5). Transcriptomic analysis revealed detectable changes in host transcriptomic homeostasis with increasing viral loads (Fig. 4), yet we did not identify any particular functional gene clusters among the differentially expressed genes. We speculate that additional factors may cooperate with Cai1 and Dicer to control viral infection, or that the tested conditions may not have reached the threshold at which elevated viral loads overcome host resilience. Further studies will address these questions.

In summary, our study reveals a complex antiviral architecture that enables cryptic mycovirus infection in the CGSC, in which mycoviruses had not been previously reported.

## Materials and Methods

The detailed description of the materials and methods used in this study is provided in the *SI Appendix*. In brief, fungal strains and culture conditions, the generation of virus-cured and gene deletion strains, total nucleic acid extraction, purification of viral particles and recombinant proteins, and quantitative RT-PCR analysis are described in detail. The *SI Appendix* further includes information on viral genome annotation, population comparative genomics and transcriptomics, RNA-seq data analysis, phylogenetic tree construction, as well as the structure prediction of the Cai1–dsRNA complex. Detailed procedures for the biotin-labeled dsRNA pull-down, biolayer interferometry (BLI) binding assays, *in vitro* endoribonuclease cleavage assays, and systematic phenotypic profiling under diverse stress conditions are also included. Information on the respective tests applied, *P* values, and numbers of replicates for each experiment is indicated in the figure captions.

## Data, Materials, and Software Availability

The nucleotide sequence of CgTV1 has been deposited in GenBank under accession number PZ721968. Metavirome sequencing data have been deposited in the NCBI BioProject database under accession number PRJNA1520860. Genomic DNA sequencing data for the 43 *C. gattii* strains were retrieved from a previously published study (33) and are available in the NCBI BioProject database under accession number PRJNA1054030. The genome assembly (FASTA file) of *C. gattii* CG1 has been deposited in GenBank-Genome under accession number JCDEVL000000000. Raw RNA-seq data for the wild-type, mutants, and virus-cured strains have been deposited in NCBI under BioProject accession number PRJNA1498692.

## Acknowledgments

This study was financially supported by the Strategic Priority Research Program of the Chinese Academy of Sciences (XDB0810000 [L.W.], [X.L.]), National Natural Science Foundation of China (32425007 [L.W.], 32522005 [X.L.]), The Chinese Academy of Sciences Project for Young Scientists in Basic Research (YSBR-111 [L.W.], [X.L.]), Beijing Natural Science Foundation (F251010 [X.L.]), and The Science & Technology Fundamental Resources Investigation Program (2022FY101100 [X.L.]). The funders had no role in study design, data collection and analysis, decision to publish, or preparation of the manuscript.

## Supporting Information

**The authors have cited additional references within the Supporting Information, which also includes supplementary figures, tables and datasets**.

**Fig. S1. RT-PCR screening of mycoviruses and validation of CgTV1 curing**.

**Fig. S2. Isolation and morphological characterization of CgTV1 virus-like particles (VLPs)**.

**Fig. S3. Volcano plot of global gene expression comparing wild-type and virus-cured *C. gattii* CG1.**

**Fig. S4. Biochemical characterization and substrate specificity of the endonuclease Cai1**.

**Fig. S5. Phenotypic assays of *C. gattii* CG1, mutant strains, and their respective virus-cured counterparts**.

**Table S1. BLASTx information of virus identified in *Cryptococcus***.

**Table S2. Wild-type, virus-cured, and engineered mutant strains of *C. gattii* used in this study. Table S3. The list of primers used in this study**.

**Table S4. Virus CP amino sequences used for pairwise alignment and phylogenetic tree. Table S5. Virus RdRp amino sequences used for pairwise alignment and phylogenetic tree**.

**Dataset S1. Information and mycovirus screening results for 386 clinical and environmental *Cryptococcus* isolates**.

**Dataset S2. Summary of 369 genes harboring high-impact mutations in *C. gattii* CG1 relative to the reference genome *C. gattii* WM276**.

**Dataset S3. List and statistical profiling of 532 candidate antiviral factors identified through *C. gattii* population transcriptomics**.

**Dataset S4. Differentially expressed genes (DEGs) in the *cai1*Δ mutant compared to its virus-cured (VC) counterpart**.

**Dataset S5. Differentially expressed genes (DEGs) in the *cai1*Δ*dcr1*Δ*dcr2*Δ mutant compared to its virus-cured (VC) counterpart**.

