## Supplementary material for "A mycovirus identified from an outbreak fungal species complex reveals a parallel antiviral system": SI Appendix; Dataset S1; Dataset S2; Dataset S3; Dataset S4; Dataset S5: Supporting Information.pdf

**This PDF file includes:**

Supporting text
Figures S1 to S5
Tables S1 to S5
Legends for Datasets S1 to S5

**Other supporting materials for this manuscript include the following:**

Datasets S1 to S5

### Supporting Information Text

### SI Materials and Methods

#### Strains and culture conditions

A total of 386 *Cryptococcus* isolates and engineered mutant strains used in this study were summarized in Dataset S1 and Table S2, respectively. Fungal cells were routinely cultured at 30°C in liquid yeast extract–peptone–dextrose (YPD) broth (1% yeast extract, 2% peptone, and 2% glucose) with shaking at 220 rpm, or maintained on solid YPD agar plates supplemented with 2% agar. For molecular cloning and plasmid propagation, *E. coli* strains were cultivated at 37°C in liquid Luria–Bertani (LB) broth with shaking at 220 rpm, or maintained on LB agar plates containing the appropriate selection antibiotics. To generate virus-cured (VC) strains, *C. gattii* CG1 cells were inoculated into liquid YPD broth supplemented with 0.4 mg/mL 2CMC and incubated with shaking at 30°C for 24 h. The treated cultures were then serially diluted and plated onto fresh YPD agar plates to obtain single colonies, and the complete elimination of CgTV1 was systematically verified by RT-PCR. Primers used for validating the VC strains were listed in Table S3.

#### Metavirome sequencing for mycovirus discovery

To screen for mycoviruses, total RNA was extracted from 386 isolates and combined into two distinct pools based on their species taxonomy: the CNSC pool and the CGSC pool. Following ribosomal RNA (rRNA) depletion, sequencing libraries were constructed and sequenced on an Illumina platform. To ensure sufficient detection power for viral transcripts within the large-scale pooled samples, deep sequencing was performed. This yielded 10.42 Gb of raw data for the CNSC pool, and 11.10 Gb of raw data for the CGSC pool.

For mycoviral discovery, raw sequencing reads were quality-filtered by Fastp (v0.22.0), and host background sequences were subtracted by mapping to the respective reference genomes *C. neoformans* H99(FungiDB-release61) and *C. gattii* WM276 (FungiDB-release68) using HISAT2 (2.2.1). The remaining unmapped reads were *de novo* assembled into contigs using rnaSPAdes (v3.15.4)(1). These contigs were subsequently queried against the NCBI non-redundant (nr) protein database using BLASTx. Contigs matching mycoviral RNA-dependent RNA polymerase (RdRp) or capsid proteins were identified as putative mycovirus sequences for further validation.

#### Nucleic acid extraction, molecular cloning, sequencing, and genomic annotation of CgTV1

We homogeneously disrupted approximately $10^8$ cells and performed total nucleic acid extraction following previously described procedures(2). To confirm the molecular identity of the viral dsRNA genome, 1 µg aliquots of total nucleic acids were independently treated with 2 U DNase I (New England Biolabs), 10 U S1 nuclease (Takara), or RNase A (Thermo Fisher) at 37°C for 1 hour. The digestion products were resolved on agarose gels to assess nuclease sensitivity.

To determine the terminal genome sequences of the CgTV1, the 5′- and 3′-terminal sequences of the viral dsRNA were characterized using a RACE kit according to the manufacturer's instructions (Vazyme Cat. No. RA101). At least three independent clones of each amplicon were sequenced in both directions by GENEWIZ (Beijing, China). Primers used for amplicon are listed in Table S3.

To characterize the genomic organization and coding potential of CgTV1, open reading frames (ORFs) were deduced using NCBI ORFfinder (https://www.ncbi.nlm.nih.gov/orffinder/). The putative –1 ribosomal frameshifting slippery site located in the overlapping junction between ORF1 and ORF2 was predicted using KnotInFrame (v2.2.0)(3).

**Phylogenetic analysis and protein similarity matrix**

Previously published sequences of *Totivirus* and closely related species were retrieved from GenBank (Table S4 and Table S5). To construct a maximum likelihood (ML) tree for the CP and RdRp independently, their amino acid sequences were aligned using MUSCLE (v5.1) with the default settings and then trimmed by trimAl (v1.4) using the automated1 option. Maximum-likelihood analyses were performed by running 1,000 bootstraps using IQ-TREE (v2.1.4) (4-6). The best-fitting substitution model of amino acid was determined by IQ-TREE (v2.1.4) and was chosen by the Bayesian information criterion. FigTree (v1.4.4; http://tree.bio.ed.ac.uk/software/figtree/) was used for visualization. Percent identity matrix (PIM) among *Totivirus* was generated by Clustal-Omega (https://www.ebi.ac.uk/Tools/msa/clustalo/).

**Purification of viral particles**

Viral particles were purified as previously described (7). $10^8$ cells were collected and ground to a fine powder in liquid nitrogen. They were mixed with 500 mM sodium phosphate buffer and centrifuged at 13,800 × *g* at 4°C for 10 min to remove cellular debris. The supernatant was then ultracentrifuged at 100,000 × *g* (OPTIMA XPN-100) at 4°C for 3 hours to collect the virus pellet, which was then resuspended in 500 mM sodium phosphate buffer. The crude virus preparation was further purified by sucrose gradient centrifugation. Subsequently aliquots of each fraction (500 μL) were subjected to dsRNA extraction to monitor for the presence of CgTV1 dsRNA. The purified virus preparations were negatively stained with 0.5% uranyl acetate solution on copper grids and then observed via transmission electron microscopy (TEM, HT7800; Hitachi). The diameter of the VLPs was measured using ImageJ (v1.54p).

**RNA purification and quantitative RT-PCR analysis**

RNA extraction and RT-qPCR were performed as previously described (8). Briefly, $10^8$ cells were collected and ground to a fine powder in liquid nitrogen. Total RNA was extracted using the Ultrapure RNA Kit (Kangweishiji, CW0581S) according to the manufacturer's instructions. Reverse transcription and RT-qPCR were performed using Fastquant Reverse Transcription kit (Tiangen KR106-02, with gDNase) and Power SYBR qPCR premix reagents (KAPA). Relative transcript levels of the CgTV1 CP gene were determined using the comparative Ct (9) method and normalized to host *ACT1* expression. Primers used for RT-qPCR are listed in Table S3.

**Population comparative genomic analysis**

Genomic sequencing data of *C. gattii* originated from previous studies (10). Raw data quality was assessed using FastQC (v0.12.1), and adapter trimming was performed using Fastp (v0.22.0). The cleaned data were mapped to the reference genome (FungiDB-release68-CgattiiWM276) using SAMtools (v1.6) and BWA-MEM (v0.7.17)(11-13). After generating VCF files using GATK HaplotypeCaller (v4.6.2.0, https://gatk.broadinstitute.org), SNPs and InDels were screened and filtered using the recommended parameters of GATK SelectVariants and GATK VariantFiltration.

Subsequently, GATK MergeVcfs was used to merge SNP and INDEL variant files. SnpEff (v5.4.0c) was utilized to annotate all variant loci and predict their functional impacts, with a focus on identifying high-impact variants(14). The mutation frequency of each gene in the population was further analyzed. Mutations in *AGO1* and *AGO2* were validated across at least three independent replicates by Sanger sequencing using the primers listed in Table S3.

**Population transcriptome analysis**

Population-level transcriptomic datasets comprising 43 *C. gattii* isolates were obtained from our previously published study (10). Within this dataset, Hierarchical Orthologous Groups (HOGs) were established across isolates using OrthoFinder (v2.5.5), with reference genome (*C. gattii* WM276) incorporated to benchmark cluster functional annotations(15). For transcriptomic profiling, raw RNA-seq reads from these 43 *C. gattii* isolates were quality-controlled using FastQC (v0.12.1) and aligned to the *C. gattii* WM276 reference genome using HISAT2 (v2.2.1)(16). Transcript abundance for each gene was quantified as transcripts per million (TPM) using StringTie (v2.1.7) and systematically mapped to its corresponding HOG(17). To screen for candidate host antiviral restriction factors specifically elevated in the mycovirus-positive isolate *C. gattii* CG1 compared to the 42 mycovirus-negative isolates, candidate HOGs were filtered using two criteria: significant population-wide divergence (top 10%, rank $\leq 4$, $Z > 1.64$, $P < 0.05$) and *C. gattii* CG1 internal baseline expression (Robust $Z > 0$).

**Structure prediction of the RNase III–dsRNA complex**

The dimeric architecture of the homologous *S. cerevisiae* Rnt1p–RNA crystal structure (PDB 5T16) suggested that Cai1 may function as a homodimer (18). Therefore, the Cai1–dsRNA complex was predicted using Protenix (v0.5.5) (19), with the full-length Cai1 sequence (293 residues) as input and the protein copy number set to two. A 33-bp dsRNA duplex (5'-GGAGAUAUACAUAUGGCGGUUGCGCAGGACUAU-3' / 5'-AUAGUCCUGCGCAACCGCCAUAUGUAUAUCUCC-3'/) was used as the experimental substrate, along with four $Mg^{2+}$. Protein multiple-sequence alignments were generated using the Protenix pipeline and enabled during inference. The resulting A3M file contained 4,222 sequence records including the query. No structural templates or additional restraints were supplied. Inference used bfloat16 precision, 10 recycling iterations, and 200 diffusion steps. Five random seeds (101–105), each generating five samples, yielded 25 candidate structures. The highest-ranking candidate free of steric clashes was selected as the representative model on the basis of the Protenix ranking_score. Model confidence was assessed by the predicted interface TM-score (ipTM), predicted TM-score (pTM), and mean predicted local distance difference test score (pLDDT).

**Construction of *C. gattii* gene deletion strains**

Gene disruption was performed as previously described (20). In brief, the 5′ and 3′ homologous fragments of the target gene were amplified via PCR and fused with a *NEO* or *NAT* resistance marker to generate a gene deletion cassette. The deletion cassette was introduced into *C. gattii* CG1 cells using the TRACE method. For the construction of complemented strains of *CAI1* in *C. gattii* CG1, the open reading frame of gene was amplified by PCR and were cloned into the plasmid pXC and introduced into the safe haven site 1 (SH1) of *C. gattii* CG1 by electroporation. For the overexpression of the *AGO1* and *AGO2* genes, we constructed plasmids containing the coding regions of these genes controlled by the *TEF1* promoter. Subsequently, these linearized plasmids were introduced into the corresponding strains by electroporation. Transformants were selected on YPD plates supplemented with neomycin (100 μg/mL), nourseothricin (50 μg/mL) or hygromycin B (200 μg/mL). All mutants were validated through PCR. Primers used for genotype verification are listed in Table S3.

**Protein purification**

The amino acid sequence of the Cai1 protein was codon-optimized using the MaxCodon™ Optimization Program (v13, DetaiBio). The target gene was synthesized *de novo* and inserted into the expression vector pET30a via the NdeI and HindIII restriction sites. Recombinant vector was confirmed by restriction digestion analysis and DNA sequencing. Subsequently, the verified plasmid was transformed into *E. coli* BL21 (DE3) strain. Cai1 protein was induced with IPTG and subsequently purified using nickel-affinity chromatography (Ni-IDA resin).

**Biolayer Interferometry Binding Assays (BLI)**

Biotin-labeled dsRNA was prepared as described above. The binding affinity between biotinylated dsRNA (23.7 nM) and purified Cai1 protein was measured by BLI on an Octet-Red 96e (Molecular Devices LLC, San Jose, CA, USA) using streptavidin-coated biosensors. The assay was performed at 37°C following a four-step sequential protocol. Briefly, biotinylated dsRNA at 23.7 nM diluted in PBS was loaded onto streptavidin biosensors. The biosensors were then immersed in PBS for 300 s to establish baseline, followed by incubation with serially diluted Cai1 protein (27–864 nM) in PBS for association, and subsequently transferred to PBS for dissociation. All data were analyzed using ForteBio Data Analysis software.

**Biotin-labeled dsRNA pull-down assay**

To synthesize biotinylated dsRNA, a DNA template harboring the T7 RNA polymerase promoter was generated by PCR using primers Pull-down-F1 and Pull-down-R2. The PCR product was purified, and its concentration was measured. Biotinylated RNA was then synthesized by in vitro transcription from the purified double-stranded DNA template. The transcription reaction mixture contained 150 ng of DNA template, 2 μL of 10 × transcription buffer, 2 μL of biotin RNA labeling mix (Roche, 11685597910), 2 μL of T7 RNA polymerase (Roche, 10881767001), and sterile RNase-free double-distilled water to a final volume of 20 μL. After gentle mixing and brief centrifugation, the reaction was incubated at 37 °C for 2 hours. To remove the DNA template, 2 μL of RNase-free DNase I was added and the mixture was incubated at 37 °C for 15 min. The reaction was stopped by adding 2 μL of 0.2 M EDTA (pH 8.0), then heated to 72 °C for 10 min and allowed to cool slowly to room temperature to promote annealing of the RNA strands into double-stranded RNA. The biotinylated RNA was purified by phenol/chloroform extraction followed by ethanol precipitation.

For RNA pull-down, Streptavidin agarose beads (SMB-B01, 150 μL) were washed with 1 mL of RNA pull-down buffer A (150 mM KCl, 25 mM Tris–HCl pH 7.4, 5 mM EDTA, 0.5% NP-40). A 1-mL protein sample (protein concentration 2–3 μg/μL) was pre-cleared by incubation with 30 μL of streptavidin beads at 4 °C for 30 min with rotation. The beads were pelleted by centrifugation at 600 × *g* for 30 s, and the supernatant was collected and diluted 2-fold with buffer A supplemented with fresh protease inhibitors. To this pre-cleared extract, 1 μg of biotinylated RNA was added and the mixture was incubated at 4 °C for 1 hour on a rotating shaker. To capture protein-RNA complexes, 30 μL of washed streptavidin agarose beads were added to each sample and incubation was continued at 4 °C for 3 hours. The beads were then washed five times with buffer A. Bound proteins were eluted by adding protein loading buffer

(10% w/v SDS, 10 mM β-mercaptoethanol, 20% v/v glycerol, 0.2 M Tris-HCl pH 6.8, and 0.05% w/v bromophenol blue) and heating at 100°C for 10 min. The eluates were resolved on pre-cast 4–12% gradient Bis-Tris gels. Protein bands of interest were visualized using a silver staining kit (ZOMANBIO, ZD303-1). The primers used for the pull-down assay are listed in Table S3.

#### *In vitro* endoribonuclease cleavage assay

To evaluate the dsRNA-cleaving activity of the candidate endoribonuclease Cai1, an *in vitro* cleavage assay was performed, alongside a commercial RNase III (Beyotime Biotechnology, Shanghai, China; Cat# R7086) as a positive control. Reactions were assembled in a total volume of 20 μL containing 1× Reaction Buffer (50 mM Tris–HCl, 50 mM NaCl, 1 mM DTT, pH 7.5), 1× $MnCl_2$ (5 mM final concentration), and 1 μg dsRNA. Three experimental conditions were set: (i) no enzyme (negative control), (ii) 0.2 U of commercial RNase III, and (iii) 40 μg of the purified nuclease. After gentle mixing, the reactions were incubated at 37 °C for 20 min. The digestion was terminated by adding EDTA to a final concentration of 50 mM (from the provided 10× EDTA solution). Aliquots (10 μL) of each reaction were mixed with 2 μL of 6× DNA loading buffer, resolved on a 1% agarose gel and visualized to evaluate substrate degradation and cleavage product profiles.

To assess the substrate specificity of Cai1, single-stranded RNA (ssRNA), single-stranded DNA (ssDNA), double-stranded RNA (dsRNA), and double-stranded DNA (dsDNA) substrates were prepared as previously described (21). Cleavage assays across these distinct nucleic acid substrates were carried out under the same reaction conditions, with an enzyme-free reaction included for each substrate to exclude non-enzymatic degradation.

#### RNA-seq analysis

For RNA-seq analysis, strains and mutants were cultured as described above. Total RNA was extracted using an Ultrapure RNA Kit (Kangweishiji; CW0581S) according to the manufacturer's instructions. RNA quality control, library construction, and high-throughput sequencing were performed on the Illumina NovaSeq X Plus platform by Tiangen Biotech (Beijing, China). RNA-seq data analysis was performed as previously described (8). Briefly, RNA-seq reads were quality controlled with Fastp (v0.22.0) and FastQC (v0.11.5). The trimmed reads were mapped to the final assembly genomes using HISAT2 (v2.2.1). The expression level for each gene was determined by StringTie (v2.1.7). DESeq2 (v1.46.0, R package) was used to calculate differentially expressed genes, significance cut-off values: | $Log_2$(fold change) | > 0.6 (~ 1.5 absolute fold change) and $P_{adj}< 0.05$ (22, 23).

#### Systematic phenotypic profiling of *C. gattii* CG1 strains and mutant strains

Susceptibility analyses to antifungal drugs and other stressors were conducted as previously described (10, 24). Each strain was incubated at 30°C in YPD liquid medium for 24 hours, then diluted to $1 \times 10^7$ CFU/mL in distilled water ($ddH_2O$) and further diluted fivefold for subsequent experiments. To assess growth phenotypes under different conditions, isolates were monitored at 39°C, or 41°C, and at pH 5.0 or pH 9.0 on YPD agar medium. For stress susceptibility analyses, prepared cells were spotted on YPD or SC medium supplemented with various stress-inducing chemicals: sorbitol (osmotic stress); glucose, mannose, trehalose, sucrose or galactose (alternative carbon sources); BSC (copper starvation); NaCl, LiCl, $CoCl_2$, $MnCl_2$, or KCl (salt or heavy-metal stress); $H_2O_2$ or menadione (oxidative stress); SDS, calcofluor white, or Congo red (cell membrane/cell wall stress); rotenone (mitochondrial inhibitors); hydroxyurea (genotoxic stress); DTT(ER/Golgi stressors); FK506 (calcineurin inhibitors); hygromycin

B or cycloheximide (translation inhibitor); caspofungin (β-glucan synthase inhibitors); Triton X-100 (membrane permeabilizing agent) ; 2CMC (inhibitor of RNA virus RdRp); antifungal agents (amphotericin B, fluconazole, 5-FC, itraconazole, terbinafine, nystatin, fludioxonil, or rapamycin) or DMSO (vehicle control). Incubation was at 30°C, and photographs were taken over 2 to 3 days.

**Statistical analysis**

All statistical analyses were performed using GraphPad Prism software (v.8.0.1). Information on the respective tests applied, *P* values, and numbers of replicates for each experiment are indicated in the figure captions. A *P*-value less than 0.05 was considered statistically significant. *$P < 0.05$; **$P < 0.01$; ***$P < 0.001$; ****$P < 0.0001$ and ns (not significant).

**Figures**

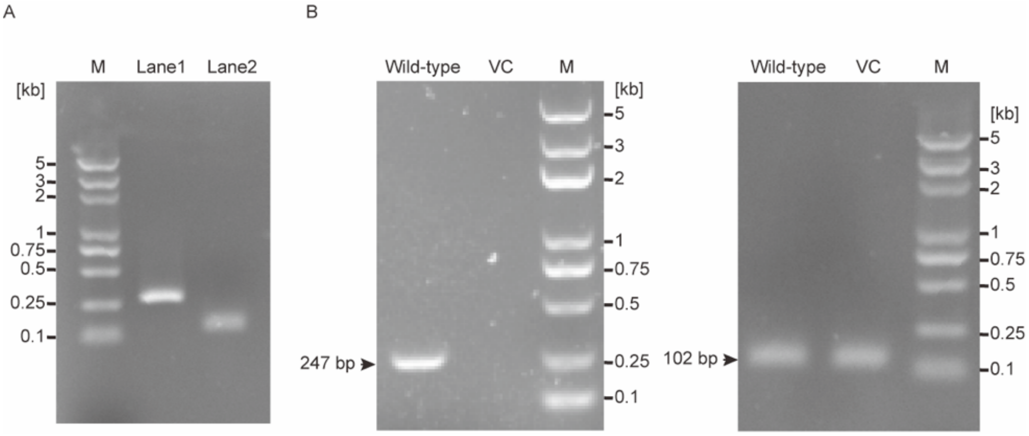


**Fig. S1. RT-PCR screening of mycoviruses and validation of CgTV1 curing.** (A) RT-PCR identification of the metavirome-discovered virus in *Cryptococcus* isolates. Among 386 screened isolates, only *C. gattii* CG1 was positive for the virus. Lane 1 shows the CgTV1-specific RT-PCR amplicon (247 bp), and Lane 2 shows the amplification of the host actin gene (*ACT1*) as an internal reference control (102 bp). (B) RT-PCR validation confirming the curing of CgTV1. The left gel demonstrates that the viral-specific amplicon (247 bp) is present in the wild-type *C. gattii* CG1 strain but completely absent in the virus-cured (VC) strain. The right gel shows the amplification of the host internal reference gene (*ACT1*, 102 bp) in both strains, verifying RNA integrity and cDNA quality. M, DNA molecular weight marker.

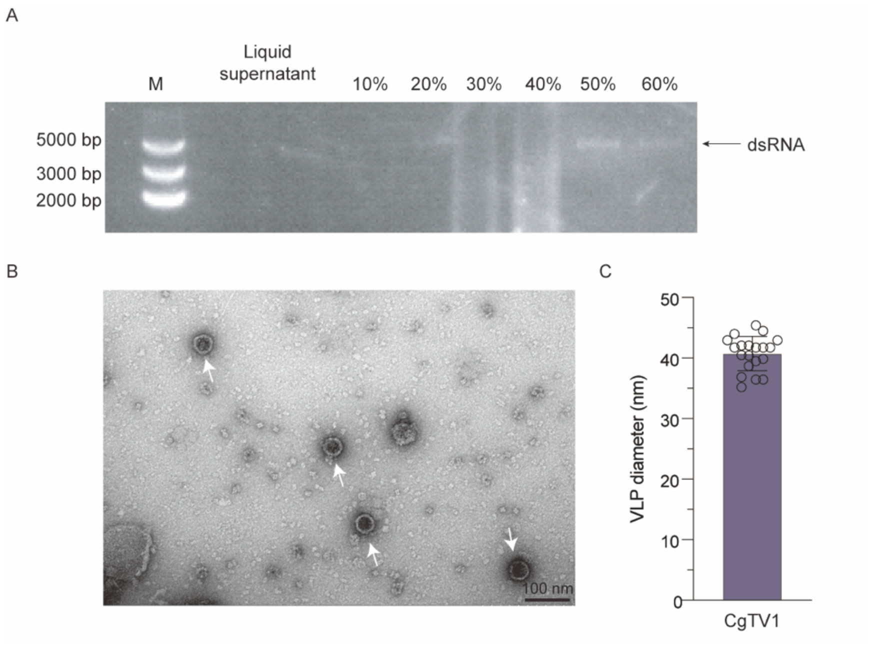


**Fig. S2. Isolation and morphological characterization of CgTV1 virus-like particles (VLPs).** (A) Agarose gel electrophoresis of dsRNA extracted from the liquid supernatant and subsequent sucrose density gradient fractions (10% to 60%, w/v). The black arrow indicates the intact CgTV1 dsRNA genome (~4.6 kb), which is predominantly enriched in the 50% and 60% sucrose fractions. M, DNA molecular weight marker. (B) Representative TEM image of negatively stained CgTV1 VLPs purified from *C. gattii* CG1. White arrows indicate intact spherical VLPs. Scale bar, 100 nm. (C) Quantitative analysis of CgTV1 VLP diameter based on TEM micrographs. Data are presented as mean ± SD (n = 20).

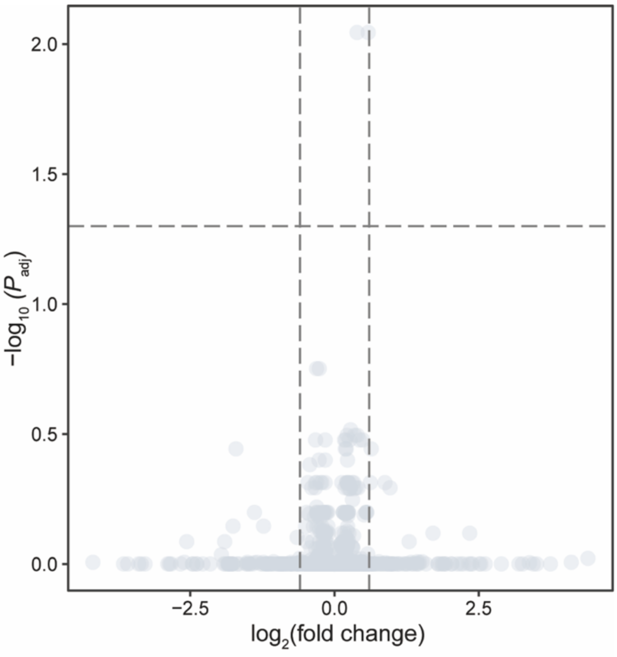


**Fig. S3. Volcano plot of global gene expression comparing wild-type and virus-cured *C. gattii* CG1.** Transcriptomic profiling reveals no DEGs meeting the combined criteria between wild-type (WT) and virus-cured (VC) strains (n = 3 biological replicates). The horizontal dashed line indicates the statistical significance threshold ($P_{adj}$ = 0.05), and the vertical dashed lines represent the fold-change cutoffs (|$\log_2$(fold change)| = 0.6, calculated as WT vs. VC). Each dot represents an individual gene. Gray dots denote non-significant genes failing to meet the combined criteria.

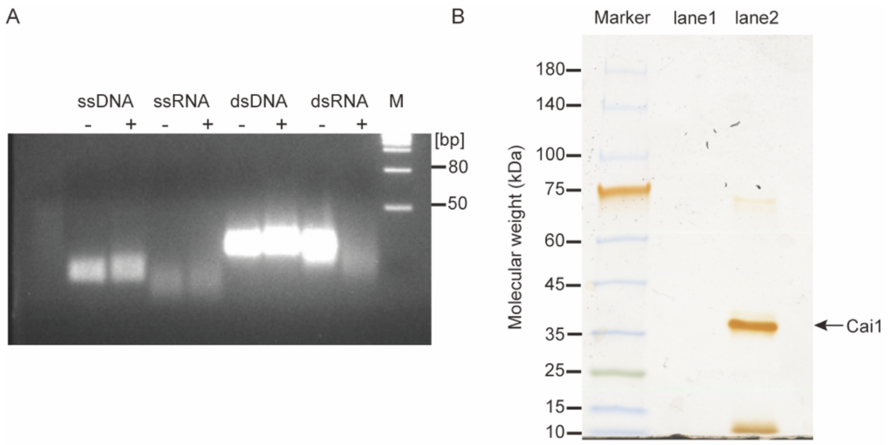


**Fig. S4. Biochemical characterization and substrate specificity of the endonuclease Cai1.** (A) *In vitro* endonuclease cleavage assays assessing the substrate specificity of recombinant Cai1. Four distinct nucleic acid substrates, namely single-stranded DNA (ssDNA), single-stranded RNA (ssRNA), double-stranded DNA (dsDNA), and native viral double-stranded RNA (dsRNA), were incubated in the absence (−) or presence (+) of purified Cai1 protein. Reaction mixtures were resolved by agarose gel electrophoresis. (B) Biotinylated dsRNA was incubated with purified Cai1 and pulled down by using streptavidin beads. The eluates were resolved by SDS-PAGE and visualized by silver staining. Cai1 was successfully pulled down in the presence of biotinylated dsRNA (lane 2), but was absent in the negative control lacking biotinylated dsRNA (lane 1). M, molecular weight markers.

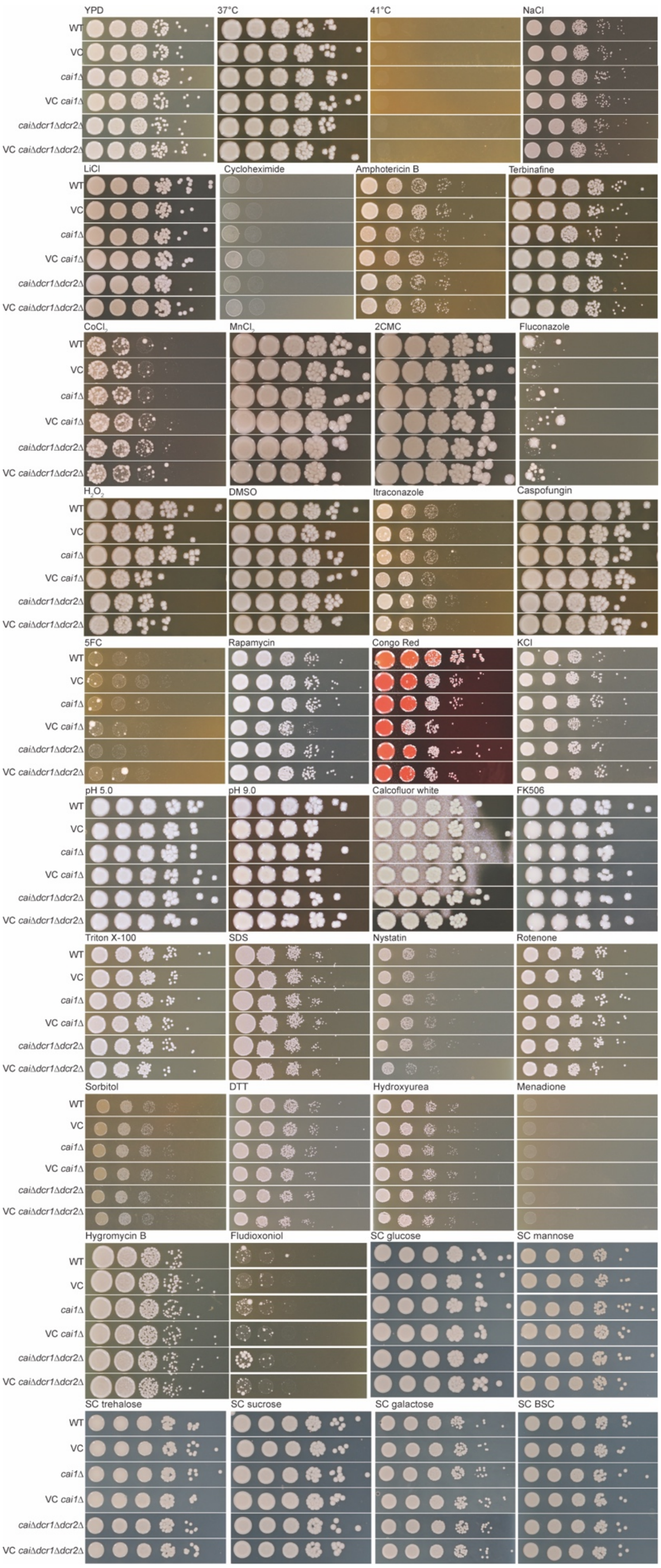


**Fig. S5. Phenotypic assays of *C. gattii* CG1, mutant strains, and their respective virus-cured counterparts.** Cells of different strains were cultured for 24 hours in YPD liquid medium at 30 °C. Cells were washed with distilled water and 5-fold serially diluted and then spotted (5 μL of each dilution) onto YPD or SC agar plates containing the indicated stress-inducing reagents. Images are representative of three independent experiments conducted with similar results.

**Tables**

**Table S1.** BLASTx information of virus identified in *Cryptococcus*.

| Contig number | Contig length (bp) | Best match | Identity | Bit-score | Genome | Genus |
|---|---|---|---|---|---|---|
| Contig 1 | 4,558 | Polyprotein (*Scheffersomyces segobiensis* virus L) | 44.90% | 723.8 | dsRNA | *Totivirus* |
| Contig 1 | 4,558 | RNA-dependent RNA polymerase, partial (*Scheffersomyces segobiensis* virus L) | 45.80% | 721.5 | dsRNA | *Totivirus* |

**Table S2.** Wild-type, virus-cured, and engineered mutant strains of *C. gattii* used in this study.

| Strain | Genotype | Source |
|---|---|---|
| *C. gattii* CG1 | Wild-type (carrying CgTV1) | This study |
| VC | CgTV1-cured wild-type derivative | This study |
| $AGO1^{OE}$ | $P_{TEF1}$-*AGO1*::*HYG* | This study |
| $AGO2^{OE}$ | $P_{TEF1}$-*AGO2*::*HYG* | This study |
| *cai1*Δ | *cai1*Δ::*NAT* | This study |
| *cai1*Δ/$P_{CAI1}$-*CAI1* | *cai1*Δ::*NAT*, $P_{CAI1}$-*CAI1*::*HYG* | This study |
| *dcr*1Δ*dcr2*Δ | *dcr*1Δ*dcr2*Δ::*NEO* | This study |
| *cai1*Δ*dcr1*Δ*dcr2*Δ | *cai1*Δ::*NAT*, *dcr1*Δ*dcr2*Δ::*NEO* | This study |

**Table S3.** The list of primers used in this study.

| Name | Sequence (5′-3′) | Purpose |
|---|---|---|
| Cas9F | GGTGACGCTGTGAGAGTGG | Cas9 amplification |
| Cas9R | GGGCCCCTCTTCACGTGG | |
| NAT-F/G418-F | GTAAAACGACGGCCAGTGC | *NAT*/*G418* amplification |
| NAT-R/G418-R | CAGGAAACAGCTATGACATGATTAC | |
| CAI1-FF | TCCTTCAAACCTGTTGCTCG | *CAI1 deletion* |
| CAI1-LF | GTCCTCCGTGACAACAGAGAATC | |
| CAI1-LR | GTCATAGCTGTTTCCTGACGAGTTTGTCGAAGGGAGG | |
| CAI1-RF | CTGGCCGTCGTTTTACTAGACGGTCCCTTCCGAG | |
| CAI1-RR | TAACGACCTCGAGAGGCCT | |
| CAI1-FR | CGCCATTCCTGGCACC | |
| CAI1-U6-R | CCACCGTAGCTGCCTGCTTCAACAGTATACCCTGCCGGTG | |
| CAI1-RNA-F | GAAGCAGGCAGCTACGGTGGGTTTTAGAGCTAGAAATAGCAAGTT | |
| dcr-FF | GGAAATGCCGACGAAGATGC | *DCR* deletion |
| dcr-LF | CGCTTTCTGATCCTGTGAAACG | |
| dcr-LR | CTGGCCGTCGTTTTACCCGGCTGATGTACGGATGAAT | |
| dcr-RF | GTCATAGCTGTTTCCTGATGGACGGAAGGAGTCGAAAG | |
| dcr-RR | TGATGGCTTACCGGGTCTTG | |
| dcr-FR | ACTGAGGGGTGCCAAACTTC | |
| dcr-U6-R | CTGTGCGCCACAATGAGAGTAACAGTATACCCTGCCGGTG | |
| dcr-RNA-F | ACTCTCATTGTGGCGCACAGGTTTTAGAGCTAGAAATAGCAAGTT | |
| NotI-$P_{CAI1}$-F | TATCCATCACACTGGCGGCCGCAGCAGAGGAAGAACAAGGCGAATGAAACGG | *CAI1* complement |
| 2417-R-PacI | TACTGCTACTGTAACCCTTAATTAATCAGACCACTTCCAGCTGCA | |
| NotI -$P_{TEF1}$-F | TATCCATCACACTGGCGGCCGCGGAAAAGGGAGGAGAAAGAAGTCAAGG | *AGO1*$^{OE}$ |
| TEF1-AGO1-R | TGGAGGCCATCATTTTGAAGTTTTCTGTGGAATCGTTAGCA | |
| AGO1-F | CTTCAAAATGATGGCCTCCAAGATCGCAG | |
| AGO1-R | TACTGCTACTGTAACCCTTAATTAATTACATGAACTATGGTGTTGCCCAT | |
| TEF1-AGO2-R | TAAGAGCCATCATTTTGAAGTTTTCTGTGGAATCGTTAGCA | *AGO2*$^{OE}$ |
| AGO2-F | CTTCAAAATGATGGCTCTTAACACTCACCCAAG | |
| AGO2-R | TACTGCTACTGTAACCCTTAATTAATTACATGAACTTACACGAATGTCAGCTCC | |
| AGO1-tF | TTTGGCTCGCCGATTTCTTCC | *AGO1* Sanger sequencing |
| AGO1-tR | GGCCATACGTGGCTGAATGTC | |

**Table S3.** The list of primers used in this study (continued).

| Name | Sequence (5′-3′) | Purpose |
|---|---|---|
| AGO2-tF | GGAGGGGAGGGAAGTGTGAAC | *AGO2* Sanger sequencing |
| AGO2-tR | TCTGATGGACAGGAAACGGC | |
| GSP1 | TGCCGGGGTCAATGCTGAATTCGT | RACE |
| GSP2 | GCAGCATGGATGGTTAAGGATGCCA | |
| CP-1 | AGCACAGATCGGAGCCAGAC | RT-PCR and qRT-PCR |
| CP-2 | CCTCTCACGAACTCAGGTAACAAC | |
| ACT-1 | GTCCTACGAGCTTCCTGACG | |
| ACT-2 | GCAGACTCGAGACCAAGGAG | |
| SH1U6-R | GCGGCTCTTCTTGTGGCTTTAACAGTATACCCTGCCGGTG | Safe haven1 sgRNA amplification |
| SH1gRNA-F | AAAGCCACAAGAAGAGCCGCGTTTTAGAGCTAGAAATAGCAAGTT | |
| Pull-down-F1 | TAATACGACTCACTATAGGGATGGCAACTACTTTGTTCGAC | pull-down |
| Pull-down-R2 | TAATACGACTCACTATAGGGTGCATATTTAGTACGGACACC | |

**Table S4.** Virus CP amino sequences used for pairwise alignment and phylogenetic tree

| CP | | | |
|---|---|---|---|
| Genus | Virus name | Abbreviation | Accession |
| *Totivirus* | *Cryptococcus gattii* totivirus 1 | CgTV1 | PZ721968 |
| | *Xanthophyllomyces dendrorhous* virus L1A | XdVL1A | YP_007697650.1 |
| | *Xanthophyllomyces dendrorhous* virus L1B | XdVL1B | YP_009507834.1 |
| | *Scheffersomyces segobiensis* virus L | SsVL | YP_009507830.1 |
| | *Tuber aestivum* virus 1 | TaV1 | YP_009507832.1 |
| | *Saccharomyces cerevisiae* virus L-BC (La) | ScVL-BC(La) | NP_042580.1 |
| | *Saccharomyces cerevisiae* virus L-A | ScVL-A | NP_620494.1 |
| *Trichomonasvirus* | *Trichomonas vaginalis* virus 1 | TvV1 | YP_009162331.1 |
| | *Trichomonas vaginalis* virus 2 | TvV2 | NP_624322.2 |
| | *Trichomonas vaginalis* virus 3 | TvV3 | NP_659389.1 |
| | *Trichomonas vaginalis* virus 4 | TvV4 | YP_009507837.1 |
| *Leishmaniavirus* | *Leishmania* RNA virus 1-1 | LrV1-1 | NP_041190.1 |
| | *Leishmania* RNA virus 2-1 | LrV2-1 | NP_043464.1 |
| *Victorivirus* | *Corynespora cassiicola* victorivirus 1 | CcVV1 | UIB81488.1 |
| | *Beauveria bassiana* victorivirus NZL/1980 | BbVV-NZL/1980 | YP_009032632.1 |
| | *Sclerotinia nivalis* victorivirus 1 | SnVV1 | YP_009259367.1 |
| | *Neofusicoccum parvum* victorivirus 3 | NpVV3 | UBZ25889.1 |
| | *Aspergillus foetidus* slow virus 1 | AfSV1 | YP_009508248.1 |
| | *Beauveria bassiana* victorivirus 1 | BbVV1 | AMQ11130.1 |
| | *Bipolaris maydis* victorivirus 1b | BmVV1b | QJW70220.1 |
| | *Nigrospora chinensis* victorivirus 1 | NcVV1 | URX65567.1 |
| | *Helminthosporium victoriae* virus 190S | HvV190S | NP_619669.2 |
| unclassified *Orthototiviridae* | *Thelebolus microsporus* totivirus 1 | TmTV1 | AZT88642.1 |
| | *Tolypocladium ophioglossoides* totivirus 1 | ToTV1 | AZT88644.1 |
| | *Rhizopus stolonifer* double-stranded RNA virus 1 | RsDSV1 | WMC21260.1 |
| | *Mucor hiemalis* virus 1 | MhV1 | CAE9672661.1 |
| *Giardiavirus* | *Giardia lamblia* virus | GLV | NP_619551.1 |
| *Chrysovirus* (outgroup) | *Helminthosporium victoriae* 145S virus | Hv145SV | YP_052859.1 |

**Table S5.** Virus RdRp amino sequences used for pairwise alignment and phylogenetic tree

| RdRP | | | |
|---|---|---|---|
| Genus | Virus name | Abbreviation | Accession |
| *Totivirus* | *Cryptococcus gattii* totivirus 1 | CgTV1 | PZ721968 |
| | *Xanthophyllomyces dendrorhous* virus L1A | XdVL1A | YP_007697651.1 |
| | *Xanthophyllomyces dendrorhous* virus L1B | XdVL1B | YP_009507835.1 |
| | *Scheffersomyces segobiensis* virus L | SsVL | YP_009507831.1 |
| | *Tuber aestivum* virus 1 | TaV1 | YP_009507833.1 |
| | *Saccharomyces cerevisiae* virus L-BC (La) | ScVL-BC(La) | NP_042581.1 |
| | *Saccharomyces cerevisiae* virus L-A | ScVL-A | NP_620495.1 |
| *Trichomonasvirus* | *Trichomonas vaginalis* virus 1 | TvV1 | YP_009162330.1 |
| | *Trichomonas vaginalis* virus 2 | TvV2 | NP_624323.2 |
| | *Trichomonas vaginalis* virus 3 | TvV3 | NP_659390.1 |
| | *Trichomonas vaginalis* virus 4 | TvV4 | YP_009507836.1 |
| *Leishmaniavirus* | *Leishmania RNA* virus 1-1 | LrV1-1 | NP_041191.1 |
| | *Leishmania* RNA virus 2-1 | LrV2-1 | NP_043465.1 |
| *Victorivirus* | *Beauveria bassiana* victorivirus NZL/1980 | BbVV-NZL/1980 | YP_009032633.1 |
| | *Corynespora cassiicola* victorivirus 1 | CcVV1 | UIB81489.1 |
| | *Sclerotinia nivalis* victorivirus 1 | SnVV1 | YP_009259368.1 |
| | *Neofusicoccum parvum* victorivirus 3 | NpVV3 | UBZ25890.1 |
| | *Aspergillus foetidus* slow virus 1 | AfSV1 | YP_009508249.1 |
| | *Beauveria bassiana* victorivirus 1 | BbVV1 | AMQ11131.1 |
| | *Bipolaris maydis* victorivirus 1b | BmVV1b | QJW70221.1 |
| | *Nigrospora chinensis* victorivirus 1 | NcVV1 | URX65568.1 |
| | *Helminthosporium victoriae* virus 190S | HvV190S | NP_619670.2 |
| unclassified *Orthototiviridae* | *Umbelopsis ramanniana* virus 2 | UrV2 | VFI65724.1 |
| | *Thelebolus microsporus* totivirus 1 | TmTV1 | AZT88643.1 |
| | *Tolypocladium ophioglossoides* totivirus 1 | ToTV1 | AZT88645.1 |
| | *Rhizopus stolonifer* double-stranded RNA virus 1 | RsDSV1 | WMC21261.1 |
| | XiangYun toti-like virus 1 | XtLV1 | UUG74250.1 |
| *Giardiavirus* | *Giardia lamblia* virus | GLV | NP_620070.1 |
| Chrysovirus (outgroup) | *Helminthosporium victoriae* 145S virus | Hv145SV | YP_052858.1 |

**Dataset S1 (separate file).** Information and mycovirus screening results for 386 clinical and environmental *Cryptococcus* isolates.

**Dataset S2 (separate file)**. Summary of 369 genes harboring high-impact mutations in *C. gattii* CG1 relative to the reference genome *C. gattii* WM276.

**Dataset S3 (separate file).** List and statistical profiling of 532 candidate antiviral factors identified through *C. gattii* population transcriptomics.

**Dataset S4 (separate file)**. Differentially expressed genes (DEGs) in the *cai1*Δ mutant compared to its virus-cured (VC) counterpart.

**Dataset S5 (separate file)**. Differentially expressed genes (DEGs) in the *cai1*Δ*dcr1*Δ*dcr2*Δ mutant compared to its virus-cured (VC) counterpart.
